# Can coastal plantation forests substitute for natural coastal forests as bird and plant habitat? A test across coastal zonation

**DOI:** 10.64898/2026.04.16.718952

**Authors:** Munehiro Kitazawa, Shin Ichikawa, Daisuke Aoki, Masayuki Senzaki

**Affiliations:** Biodiversity Division, National Institute for Environmental Studies, Tsukuba, Ibaraki 305-8506, Japan; Graduate School of Agriculture, Hokkaido University, Sapporo, Hokkaido 060-8589, Japan; Japanese Migratory Bird Research Group, Faculty of Environmental Earth Science, Hokkaido University, Sapporo, Hokkaido 060-0810, Japan; Department of Wildlife Biology, Forestry and Forest Products Research Institute, Tsukuba, Ibaraki 305-8687, Japan; Faculty of Environmental Earth Science, Hokkaido University, Sapporo, Hokkaido 060-0810, Japan

**Keywords:** Biodiversity conservation, Brown shrike, Coastal forest, Natural forest, Plantation forest, Sand-dune grassland, Stand age

## Abstract

Natural forests are increasingly being replaced by plantation forests, highlighting an urgent need to reconcile forest use and biodiversity conservation. Coastal forests serve as refugia for species of conservation concern, but they have been globally replaced by plantations, and conversion remains ongoing. As natural coastal forests become increasingly scarce, opportunities for direct biodiversity comparisons with coastal plantations are also disappearing. This scarcity leads to underestimation of the conservation value of natural coastal forests, while effective management measures to conserve biodiversity within coastal plantations remain poorly developed. By focusing on the largest natural coastal forests in Japan, we assessed whether plantation coastal forests can substitute for natural coastal habitats across multiple taxonomic groups, functional groups, and threatened species. We found that plantations supported bird species richness comparable to natural forests at the community level and for threatened species. Surveyed plantation forests supported 7% of global breeding populations of the endangered Brown Shrike. However, plantations hosted lower forest plant species and higher introduced plant species richness. Plantations also significantly altered coastal zonation patterns in plant communities: forest species distribution shifted seaward, whereas coastal and introduced species shifted landward. Coastal plantations are unlikely to be complete substitutes for natural coastal forests for plant species, yet still have the potential for restoring historically lost coastal forest ecosystems. Confining further plantation activities to landward areas and plantations with low canopy cover will mitigate the negative impacts of plantations and further conserve unique coastal communities where forest and grassland species can coexist.

**Highlight:**

✓ Coastal forests have been historically disturbed and converted to plantations
✓ We test whether coastal plantations can substitute for natural forests across taxa
✓ Bird species richness in plantations was comparable to that in natural forests
✓ Plant species richness was lower in plantations and zonation patterns were altered
✓ Considering zonation patterns can turn plantations into restoration opportunities

## 1. Introduction

Humans have severely impacted the biosphere worldwide, resulting in current biodiversity crisis. The most significant direct driver is land-cover changes, such as massive clearance of natural forests (Jaureguiberry *et al*. 2022; Maxwell *et al*. 2016). We are still facing the unprecedented rates of natural forest losses (Curtis *et al*. 2018; Watson *et al*. 2018; Winkler *et al*. 2021), and thus protecting remaining natural forests and retaining their integrity represents a central conservation priority in forest ecosystems (Grantham *et al*. 2020). The other promising pathway is to restore and improve already modified forests (Lindenmayer *et al*. 2012), as the area of plantation forest is rapidly increasing (Curtis *et al*. 2018; Mishra *et al*. 2022). Therefore, quantifying the extent to which plantation forests can substitute for natural forest habitats and the degree to which their habitats can be enhanced through management provides fundamental knowledge to reconcile forestry and biodiversity conservation.

Biodiversity responses to plantations can vary widely with forest types and stand ages (Hua *et al*. 2022; Wang *et al*. 2022). Plantation forests as habitat for wildlife species have thus been compared with natural forests across a wide range of forest types, including secondary and restored forest (Barlow *et al*. 2007; Hua *et al*. 2022), riparian forest (Martín-García *et al*. 2013), savanna forest (Didas *et al*. 2022), tropical peatland forest (Warren-Thomas *et al*. 2022), and mangrove forest (Su *et al*. 2021) to inform biodiversity conservation within plantations. Among these, coastal forest remains one of the major forest types that are still understudied and thus represent a key missing piece in comprehensively understanding the extent to which plantations can substitute for natural forest habitats (Bonari *et al*. 2017; Muñoz-Reinoso 2021).

Over centuries, natural coastal forests have been extensively converted to plantations worldwide, from the Atlantic, Baltic, and Mediterranean coasts (Muñoz-Reinoso 2021; Provoost *et al*. 2011; Reisch & Hartig 2021) to East Asia, including Japan and South Korea (Muñoz-Reinoso 2021; Nagamatsu & Matsushima 2014). These suggest that humans have almost completed the conversion of natural coastal forests to plantations and continue to expand plantations, even though we have a very limited understanding of the ecological consequences of coastal forest plantations. Given that coastal ecosystems can provide refugia for species of conservation concern (Acosta *et al*. 2009; Martínez & Psuty 2007; Sperandii *et al*. 2021), identifying practical management interventions that curb biodiversity loss in coastal ecosystems is a pressing priority.

To bridge this gap, we evaluated the extent to which coastal plantation forests can substitute for natural coastal habitats across multiple taxonomic groups, functional groups, and species spanning different extinction-risk categories. Coastal ecosystems are composed of coastal forest and sand-dune grassland, and well-preserved coastal ecosystems are characterized by zonation patterns (Acosta *et al*. 2007; Torca *et al*. 2019) where the dominant functional group shifts from the shoreline to the backshore along disturbance gradients such as salt spray, wind, and sandy sediments (Barbour & DeJong 1977; Hesp 1991; Martínez & Psuty 2007; Maun & Perumal 1999; Shimamura *et al*. 2006). Wind fences are often installed together with plantations to promote the growth of planted trees, which may disrupt gradients in coastal stresses and thereby alter zonation patterns in coastal species richness at both the community and functional-group levels (Fig. 1B). Therefore, in highly dynamic coastal ecosystems, evaluating plantation impacts across multiple co-occurring functional groups is essential for assessing potential shifts in community composition across management types and distance from the shoreline. Although stand age dictates species richness in plantation forests (Hua *et al*. 2022), this relationship has rarely been assessed in plantation coastal forests.

**Fig. 1.**
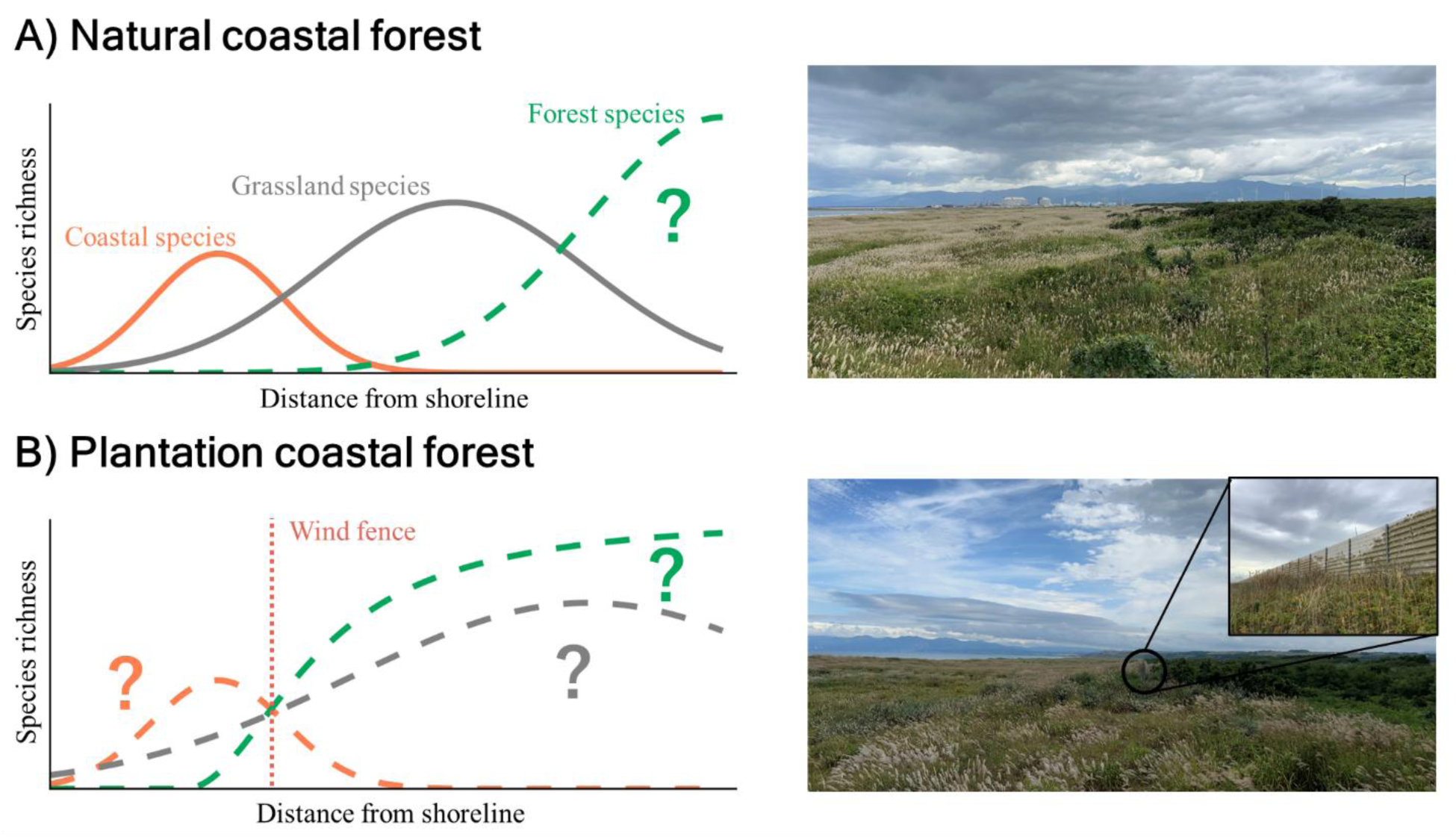
Conceptual framework of this study. A) Predicted species richness zonation patterns in natural coast and the photograph of natural sand-dune grassland and coastal forest along the Ishikari coast. Dominant functional groups are expected to shift from coastal to grassland to forest species with increasing distance from shoreline, but the extent to which plantations can substitute for natural habitats remains unclear. B) Potential species richness zonation patterns in planted coast and the photographs of natural sand-dune grassland and coastal plantation forest along the Ishikari coast. Coastal plantation forests are often characterized by tall and massive windbreaks that can disrupt species richness zonation patterns.

Here, our purpose is threefold. First, we directly compared bird and plant species richness among sand dune grasslands, natural coastal forests, and coastal plantation forests at the community and functional-group level. Effects of plantation and subsequent management are often taxon-specific (Duflot *et al*. 2025; Irwin *et al*. 2014), highlighting the need to assess and consider multiple taxa simultaneously when designing forest management plans. Second, we assessed whether plantations alter zonation patterns along the distance from the shoreline by comparing shifts in community- and functional group-level species richness between the natural and plantation coast. We further evaluated how forest structure and zonation patterns shape the occurrence of endangered bird species, thereby providing evidence-based information for local conservation and management. Third, we quantified whether stand age remains an important driver explaining forest biodiversity in plantation coastal forests by assessing the relationships between stand age and bird and plant species richness. Plant and avian field surveys were conducted along the Ishikari coast in northern Japan, where natural coastal forests are well preserved (Matsushima *et al*. 2014), which provides a rare opportunity to test whether coastal plantation forests can substitute for natural coastal forests.

## 2. Materials and Methods

### 2.1. Study area

Field surveys were carried out along the Ishikari coast, northern Japan (43°09′–43°17′ N, 141°13′ –141°23′ E, mean annual temperature: 7.9 °C, mean annual precipitation: 994 mm). Forests and wetlands in the Ishikari lowland remained largely undisturbed until around 1880 (Kitazawa *et al*. 2022b), and coastal forests within 1 km of the shoreline have been protected by the local government since 1893 (Hasegawa 1984). Therefore, vegetation zonation from bare sand through sand-dune grassland and shrub to coastal forest is well preserved along the Ishikari coast (Matsushima et al., 2014). Sand-dune grasslands were dominated by *Carex microtricha*, *Miscanthus sinensis*, *Rosa rugosa*, *Dianthus superbus*, whereas the seaward natural coastal forests consisted mainly of pure stands of *Quercus dentata* (Fig. S1) (Hasegawa 1984; Ishikari Beach Coastal Plant Conservation Center 2006). *Quercus dentata* in this region regenerates primarily through sprouting, allowing for pure stands in highly disturbed coastal forests (Hasegawa 1984). Some part of sand-dune grasslands and coastal forests had been converted to coastal plantations, which were dominated by planted *Quercus dentata* (Fig. S1), with some stands composed of introduced *Robinia pseudoacacia* or *Populus alba*.

### 2.2. Sample plot selection

We classified forest stands along the Ishikari Coast as natural or plantation using forest inventory records provided by the local government. All grasslands on the seaward side of the coastal forest, including bare sand with sparse vegetation, were classified as sand-dune grassland. Most sand-dune grasslands had experienced little direct human modification, and port areas, landfill areas, and heavily transformed grasslands were excluded when establishing bird survey plots and vegetation transects. We then established four, 16, and 10 bird survey plots for sand-dune grasslands, natural coastal forests, and plantation coastal forests, respectively. All bird sample plots were 150 m in length and 100 m in width (1.5 ha). In each plot, we established a survey line running parallel to the shoreline through the plot center (Fig. S2). Plots were randomly placed along the Ishikari Coast and spaced at least 150 m apart. Because this spacing may not have been sufficient to fully exclude repeated records of the same individual at neighboring plots (Hanioka *et al*. 2018), we moved between adjacent plots while continuously checking for birds moving between survey locations. We did not observe any such movements during the surveys, which should have reduced the possibilities of repeated records from individuals using nearby plots. Bird sample plots were distributed from 50 to 770 m from the shoreline, covering habitats from bare sand to the landward natural coastal forests.

We then laid out 15 vegetation survey lines perpendicular to the avian survey lines. All vegetation survey lines were 300 m long, extending from the shoreline to the seaward coastal forests to encompass major shifts in dominant plant functional groups (Fig. S2). We established 11 vegetation quadrats (4×4 m) along each vegetation survey line at 30 m intervals, yielding a total of 165 quadrats. Six vegetation quadrats were excluded from our surveys because they fell within residential areas. As a result, we surveyed 105, 21, and 33 vegetation quadrats for sand-dune grasslands, natural coastal forests, and plantation coastal forests, respectively.

### 2.3. Bird and plant surveys

We performed bird community surveys in each sample plot throughout the breeding season (May–July) of 2022. Each plot was surveyed three times on different days. A single surveyor (M.K.) slowly walked the survey lines (2 km/h) using a global positioning system device and recorded the species, sex, location, and behavior (e.g., singing, territorial conflict) of all detected individuals within plots (Bibby *et al*. 2000). Surveys were completed from dawn to 0900 and avoided rain and strong wind (wind speed<4 m/s). We used the maximum number of observed individuals (i.e., number of territories) for each species across the three surveys as an abundance metric. We used the summed abundances across species and the number of species detected across three surveys as response variables at the community and functional-group levels in the following analyses. We classified all observed species into one of two functional groups (i.e., forest or openland species) based on the JAVIAN database (Takagawa *et al*. 2011). In this study, we pooled grassland, wetland, and bare-ground species, which were treated separately in the previous study (Kitazawa *et al*. 2022b), into openland species because abundances in each group were low.

We also surveyed vegetation once in each vegetation quadrat in August 2022. At each quadrat, a single observer (M.K.) identified all vascular plant species and visually estimated their percentage cover in 10% classes. Plant species richness was quantified as the total number of species recorded per quadrat. We recorded both native and planted species within the quadrat.

We classified all observed plant species into one of four functional groups (i.e., coastal, grassland, forest, or introduced species) based on (Ohashi *et al*. 2015). We also calculated wood cover within each avian survey plot using aerial photographs captured in August 2020, published by the Geospatial Information Authority of Japan (https://maps.gsi.go.jp/). In addition, we scanned each avian survey plot with the SiteScape app (https://www.sitescape.ai/) on an iPhone 13 Pro to construct 3D models of the environment, from which we extracted maximum vegetation height. After manually removing noise points using CloudCompare (https://www.cloudcompare.org/), we estimated maximum vegetation height for each plot as the vertical difference between the uppermost vegetation points and the underlying ground points.

### 2.4. Statistical analysis

We conducted three sets of analyses separately for avian and plant communities. For each taxon, we (i) compared species richness between plantation and natural coastal forest, (ii) assessed whether plantations alter zonation patterns, and (iii) quantified the effects of stand age on species richness in plantation forests. These analyses were conducted primarily using generalized linear mixed models (GLMMs) in R ver. 4.5.1 (R core team 2025) and the lme4 package ver.1.1.37 (Bates *et al*. 2015). Avian plot ID and vegetation survey line ID were included as a random intercept in the avian and plant models, respectively, to account for unmeasured site-specific heterogeneity. All continuous explanatory variables were standardized prior to the analysis. The detection process of avian species was not considered in the model because openland songbirds have relatively high detectability (∼ 0.66) (Yamaura *et al*. 2016), and the detectability of birds’ calls within 50 m from the observer is high and stable among different habitats (Schieck 1997). We calculated 95% confidence intervals for model coefficients and considered effects to be statistically significant when the interval did not overlap zero or among land-cover categories.

#### 2.4.1. Species richness in grasslands, coastal natural forests, and plantation forests

We first compared community- and functional group-level bird and plant species richness among sand-dune grasslands, natural coastal forests, and plantation coastal forests using GLMMs with a Poisson error distribution and a log link function. Separate models were fitted for each taxon and level of ecological organization (community or functional-group levels), with species richness as the response variable. Land cover category (i.e., sand-dune grasslands, natural coastal forests, or plantation coastal forests) was included as the sole fixed effect, and we omitted the intercept so that model coefficients directly represented expected species richness in each land-cover category (Kéry 2010).

#### 2.4.2. Bird zonation patterns

We then assessed whether plantations alter zonation patterns in coastal ecosystems. We expected a linear zonation pattern in bird species richness along the distance from the shoreline because we focused on two contrasting bird functional groups: forest species and openland species. The former is expected to prefer more stable landward habitats, whereas the latter is associated with open habitats maintained by coastal disturbance. We modelled community- and functional group-level responses using GLMMs with a Poisson error distribution and a log link function, with bird species richness or abundance as the response variable. Land cover category (i.e., natural or plantation: 0 or 1), distance from shoreline (m), and the interaction term between these two variables were included as explanatory variables. We also fitted an alternative GLMM including a quadratic term for distance from the shoreline. This model had higher AIC values (community: 162.4; grassland species: 118.2) than the linear model (community: 161.5; grassland species: 117.6), except for forest species. Therefore, we used a linear model for bird analysis.

We next assessed whether coastal plantations alter zonation patterns at the species level for both common and endangered openland avian species. Threatened species were identified using the fourth national Red List and the revised version of Hokkaido Red List. (https://jpnrdb.com/).

Because all threatened species detected during our surveys were open-land species, subsequent analyses were restricted to this group. We excluded species recorded in fewer than five plots to avoid unstable estimates. None of the excluded species was listed in either Red List. Separate generalized linear mixed models (GLMMs) with a Poisson error distribution and a log link function were fitted for each species, with species-level abundance as the response variable.

Land cover category (i.e., natural or plantation: 0 or 1), distance from shoreline (m), and the interaction term between these two variables were included as explanatory variables. In species-level analyses, we did not include a quadratic term for distance because detections were dominated by zeros and the linear and quadratic distance terms showed strong multicollinearity, with VIF values exceeding 25 in the Brown Shrike model.

To further identify the effect of vegetation structure underlying the species-level bird abundance patterns along the distance from the shoreline, we established two additional models. First, to quantify the effect of wood vegetation structure, we fitted a Poisson GLMM with a log link function that included wood cover within the plot as linear and quadratic terms, together with vegetation height, as fixed effects. Because multicollinearity between the linear and quadratic wood-cover terms was low, with VIF values below 4 in the Brown Shrike model, we retained the quadratic term in this model. For the second model, we used a generalized linear model (GLM), which included stand age as the sole fixed effect. We used GLMs instead of GLMMs because the number of detection events for individual species was too sparse to predict random effects.

#### 2.4.3. Plant zonation patterns

For plants, we expected a unimodal pattern in species richness along the distance from the shoreline because we focused on multiple functional groups that are expected to respond differently along a gradient of coastal disturbance frequencies. Separate models were fitted at the community and functional-group levels, with plant species richness as the response variable. Land-cover category (natural vs plantation), distance from the shoreline, the quadratic term for distance, and the interaction between land-cover category and distance were included as fixed effects, and vegetation transect ID was included as a random intercept. We included the interaction term to evaluate whether plantations modify plant species richness peaks along distance from the shoreline. Positive interaction coefficients indicate greater distances from the shoreline in plantations, in other words, an inland shift in the species richness peak in plantations. We also constructed an alternative model using a dataset excluding planted tree species; however, the results were qualitatively similar to those of the main analysis (Fig. S3). In both the avian and plant models, we pooled sand-dune grassland and natural coastal forest plots into a single “natural vegetation” category because these habitats were not disturbed by plantation activities.

#### 2.4.4. Stand age

Finally, we quantified the effects of stand age on community- and functional group-level bird and plant species richness in plantation forests. We used a GLMM with a Poisson error distribution and log link. Species richness was used as the response variable, and stand age obtained from forest inventory data was included as the sole fixed effect.

## 3. Results

### 3.1. Species richness in grasslands, natural coastal forests, and plantation forests

We recorded a total of 338 individuals of 34 bird species (openland species: 173 individuals of 14 species; forest species: 165 individuals of 20 species) and 86 vascular plant species (13, 38, 14, and 21 species for coastal, grassland, forest, and introduced species, respectively) (Table S1). We detected two endangered openland birds, including the Chestnut-eared Bunting *Emberiza fucata,* and 12 territories of the Brown Shrike *Lanius cristatus superciliosus*. The number of Brown Shrikes tally alone corresponds to roughly 8% of the estimated global breeding population for this endemic subspecies (Kitazawa *et al*. 2022a).

At the community level, species richness in plantation coastal forests was comparable to that in sand-dune grassland and natural coastal forests for both birds and vascular plants (Fig. 2A). At the functional group level, both forest and openland bird species richness in natural forests were comparable to those in plantation forests (Fig. 2B). Forest bird species richness was significantly higher in natural forests than in grassland (Fig. 2B). However, plant functional group-level species richness differed among land-cover types, indicating contrasting responses to plantation among functional groups. Coastal plant species richness was highest in grasslands, intermediate in plantation forests, and lowest in natural forest (Fig. 2C). For grassland plant, species richness in natural forests was comparable to that in grassland but was significantly lower than that in plantation forests (Fig. 2C). Forest plant species richness was by far highest in natural forests among all land-cover types (Fig. 2C). Species richness of introduced plants was highest in plantation forests (Fig. 2C).

**Fig. 2.**
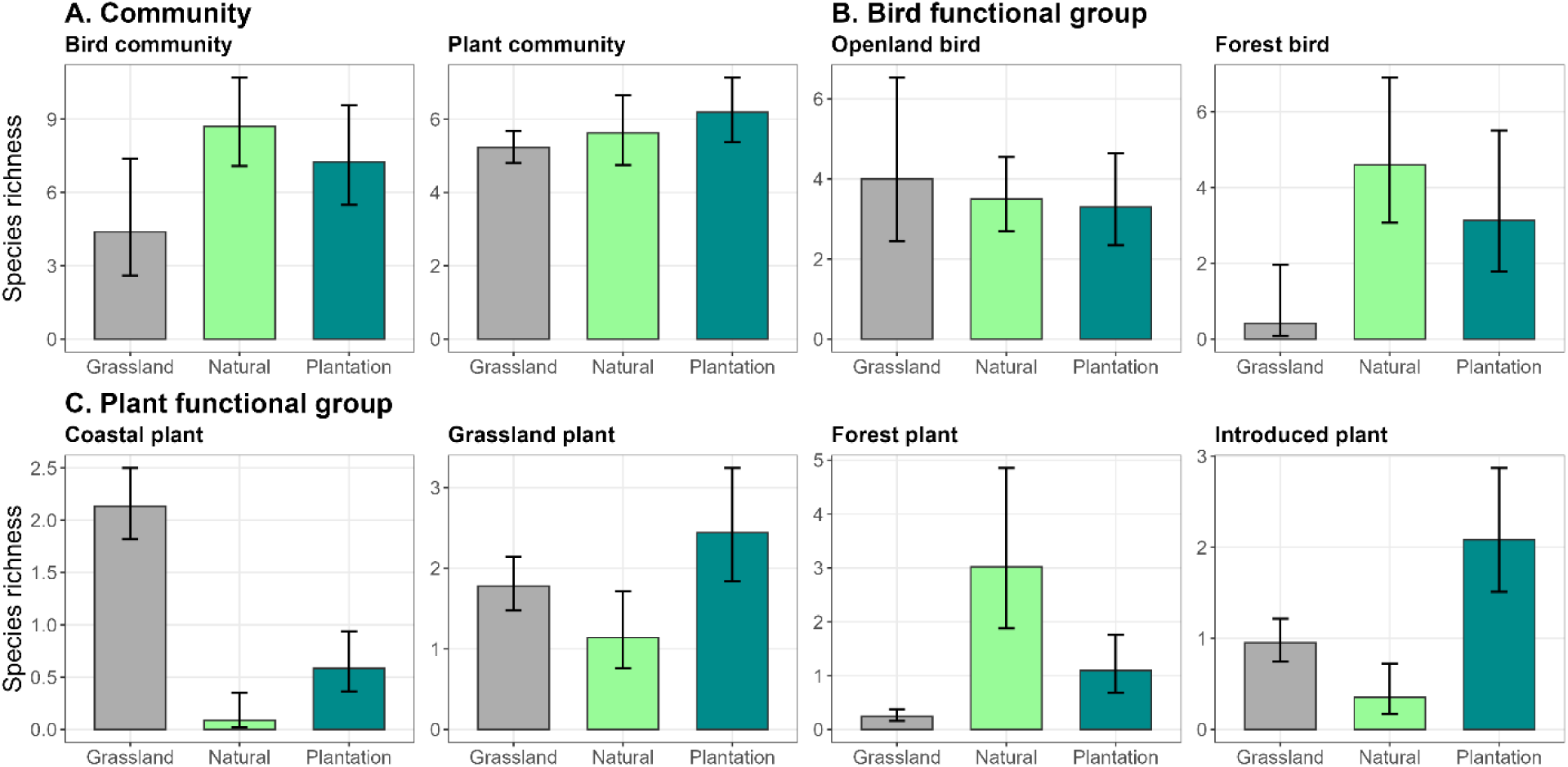
Community- and functional group-level species richness among sand-dune grasslands, natural coastal forests, and plantation coastal forests for avian (upper row) and plant species (middle and lower row). Error bars indicate 95% confident intervals. Bird species richness was calculated per 1.5 ha, and plant species richness per 16 m^2^.

### 3.2. Zonation patterns

After controlling for distance from shoreline, bird species richness in plantation forests was still comparable to that in natural vegetation at both the community and functional group levels (Fig. 3a, d, g). The effects of distance from shoreline had significantly positive effects on bird community- and forest-group species richness, and significantly negative effects on openland species richness (Fig. 3a-i). The interaction terms between plantation category and distance were not significant across all avian models (Fig. 3a, d, g), suggesting that plantation did not significantly alter coastal zonation patterns in bird species richness. At the species level, the abundances of all analyzed openland species in plantations were comparable to those in natural vegetation (Fig. 4, S4-S8), although three of the six species showed significantly higher abundances in younger plantation forests. The quadratic term for tree cover had significant effects on three of the six openland species, including the endangered Brown Shrike and Chestnut-eared Bunting, indicating that those species preferred habitats composed of a mixture of grassland and wood cover.

**Fig. 3.**
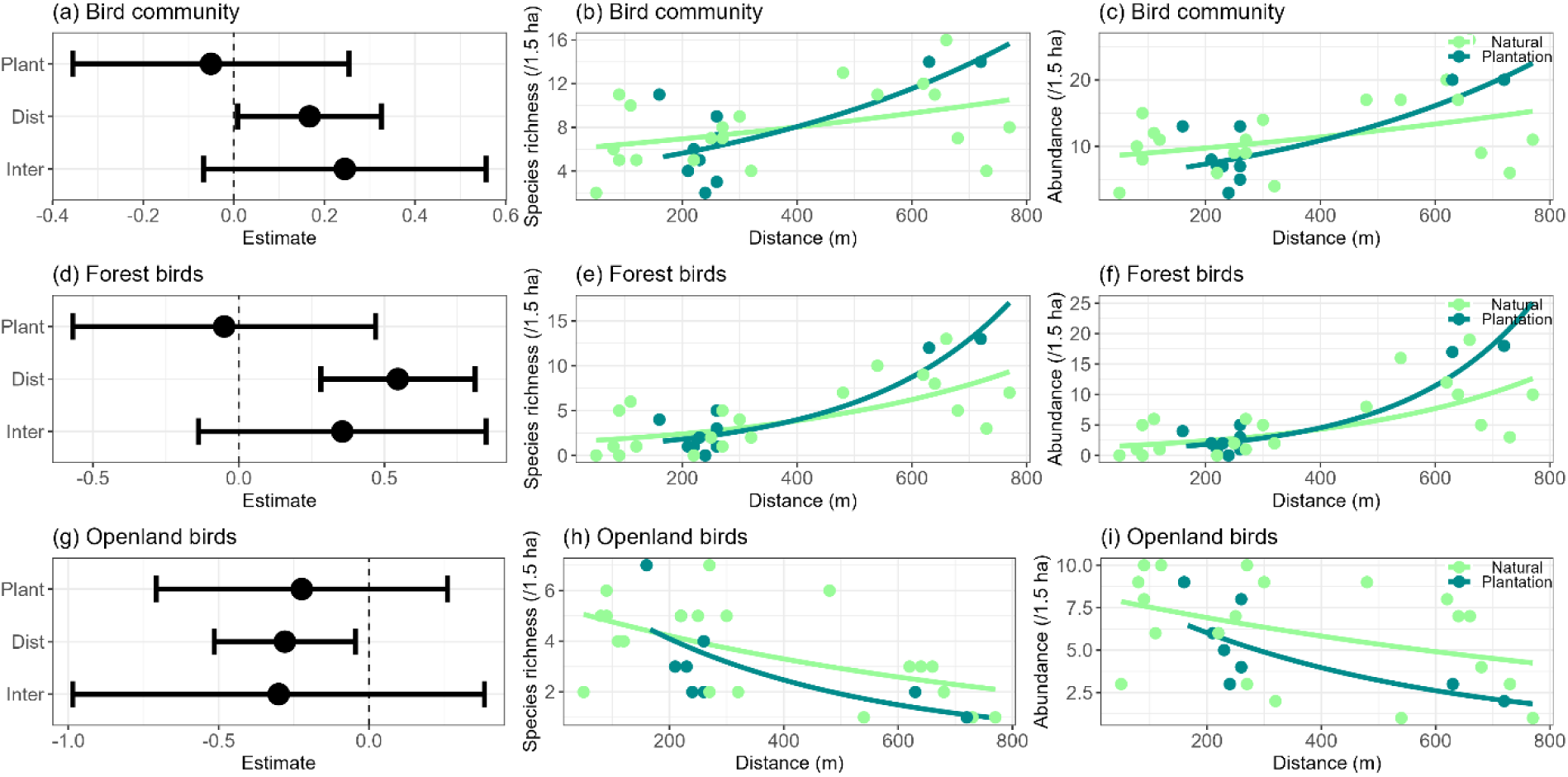
Relationships between bird community-level (a-c) and functional group-level (d-i) species richness and plantation category (Plant), distance from the shoreline (Dista), and their interactions (Inter). Left-hand panels show estimated coefficients (dots) with 95% confidence intervals (error bars) for species richness model. Middle and right-hand panels show predicted relationships between distance from shoreline and species richness and abundance, respectively. Lines represent model predictions, and dots indicate observed values for natural forests (light green) and plantation forests (dark green).

**Fig. 4.**
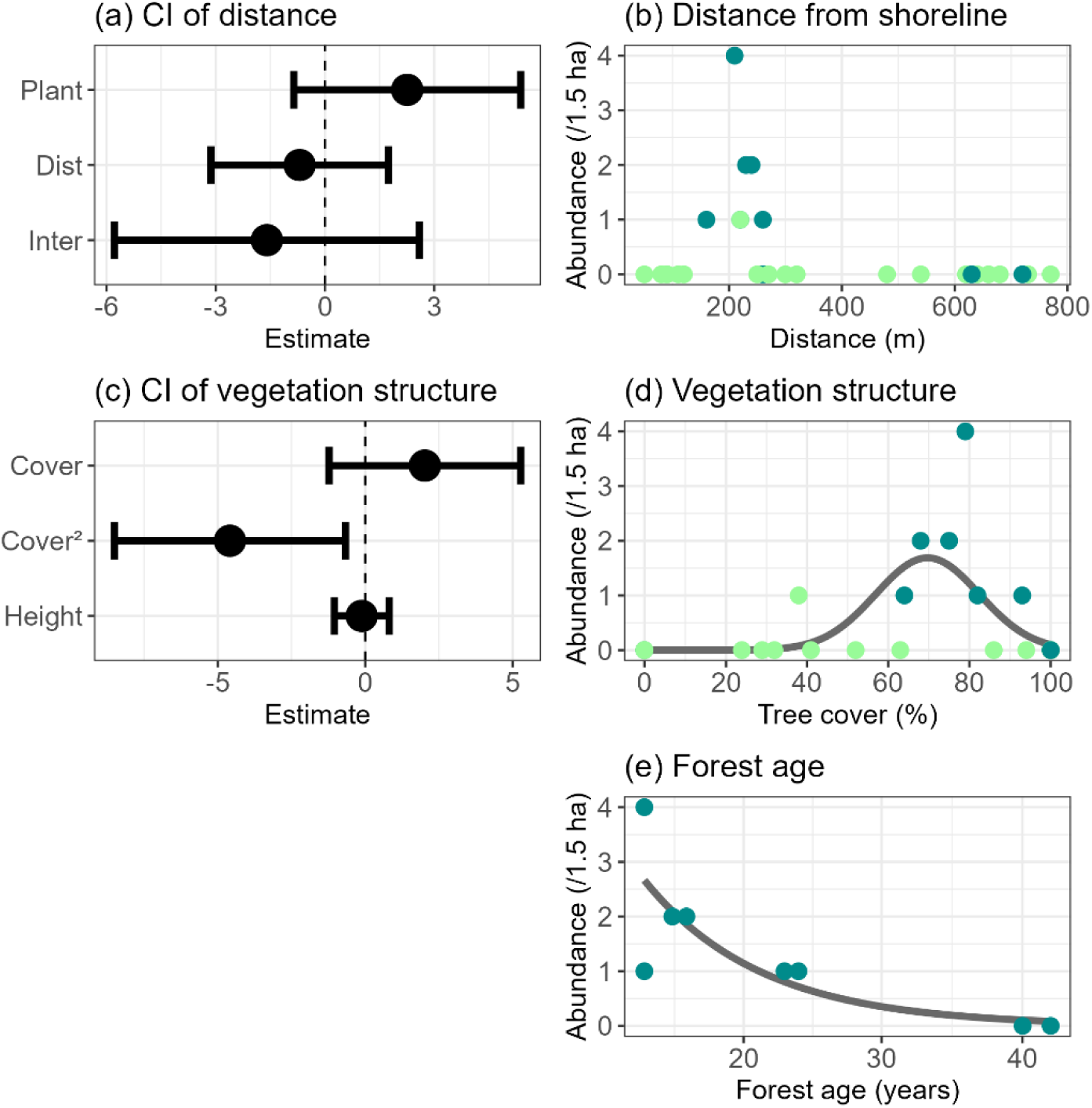
Relationships between endangered Brown Shrike abundance and environmental variables. (a)-(b) Estimated coefficients and confident intervals for plantation category (Plant), distance from the shoreline (Dista), and their interactions (Inter). (c)-(d) Estimated coefficients and confident intervals for wood cover (Cover) and its squared term (Cover^2^), and max vegetation height (Height). (e) Predicted relationships between Brown Shrike abundance and stand age. Left-hand panels show estimated coefficients (dots) with 95% confidence intervals (error bars) for each predictor. Right-hand panels show predicted relationships between (b) distance from shoreline, (d) tree cover, and (e) stand age and Brown Shrike abundance, respectively. Lines represent model predictions, and dots indicate observed values for natural forests (light green) and plantation forests (dark green). Since the global population size of this species is fairly low, predicted habitat preferences may be biased by density-dependent habitat selection, whereby individuals preferentially use sites with relatively high conspecific density.

Community-level plant species richness was significantly lower in plantations than in natural vegetation after controlling for distance from the shoreline (Fig. 5a). The interaction between plantation category and distance was not statistically significant (Fig. 5a, b), indicating that plantations still retained zonation patterns in community-level species richness. In contrast, at the functional group-level, interaction terms for coastal, forest, and introduced species had significant effects on species richness (Fig. 5c-j), suggesting that plantations altered zonation patterns in community compositions in coastal ecosystems. Peaks in coastal and introduced plant species richness were shifted landward in plantation forests (Fig. 5d, i), whereas the peak in forest plant species richness shifted seaward (Fig. 5h).

**Fig. 5.**
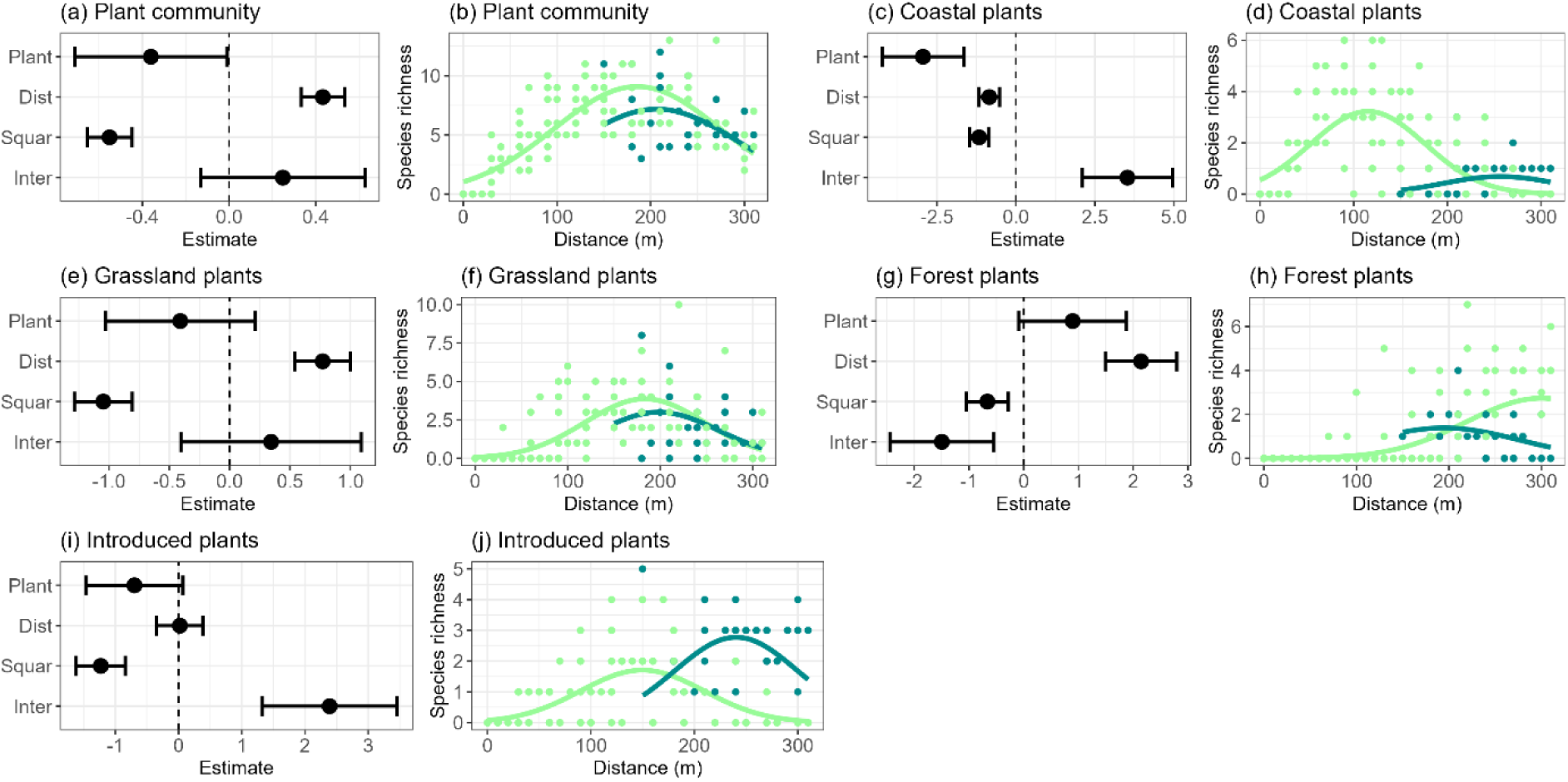
Relationships between plant community-level (a-b) and functional group-level (c-j) species richness and plantation category (Plant), distance from the shoreline (Dista), its squared term (Squar), and interaction between plantation category and distance (Inter). Left-hand panel for each group shows estimated coefficients (dots) with 95% confidence intervals (error bars) for each predictor. Right-hand panels for each group show predicted relationships between distance from shoreline and species richness. Lines represent model predictions, and dots indicate observed values for natural forests (light green) and plantation forests (dark green).

### 3.3. Stand age

Stand age in surveyed plantation forests ranged from 13 to 61 years since plantation. Two plantation bird survey plots were 61 years old, but most canopy tree had died back there (Fig. S9). We therefore excluded these plots from the main analysis. Stand age had significant positive effects on bird community and forest species richness (Fig. 6). No significant effects were detected for openland bird species richness and plant species richness, which may reflect limited sample size and restricted variation in stand age. An additional analysis, including the dieback plots, yielded qualitatively similar results (Fig. S10).

**Fig. 6.**
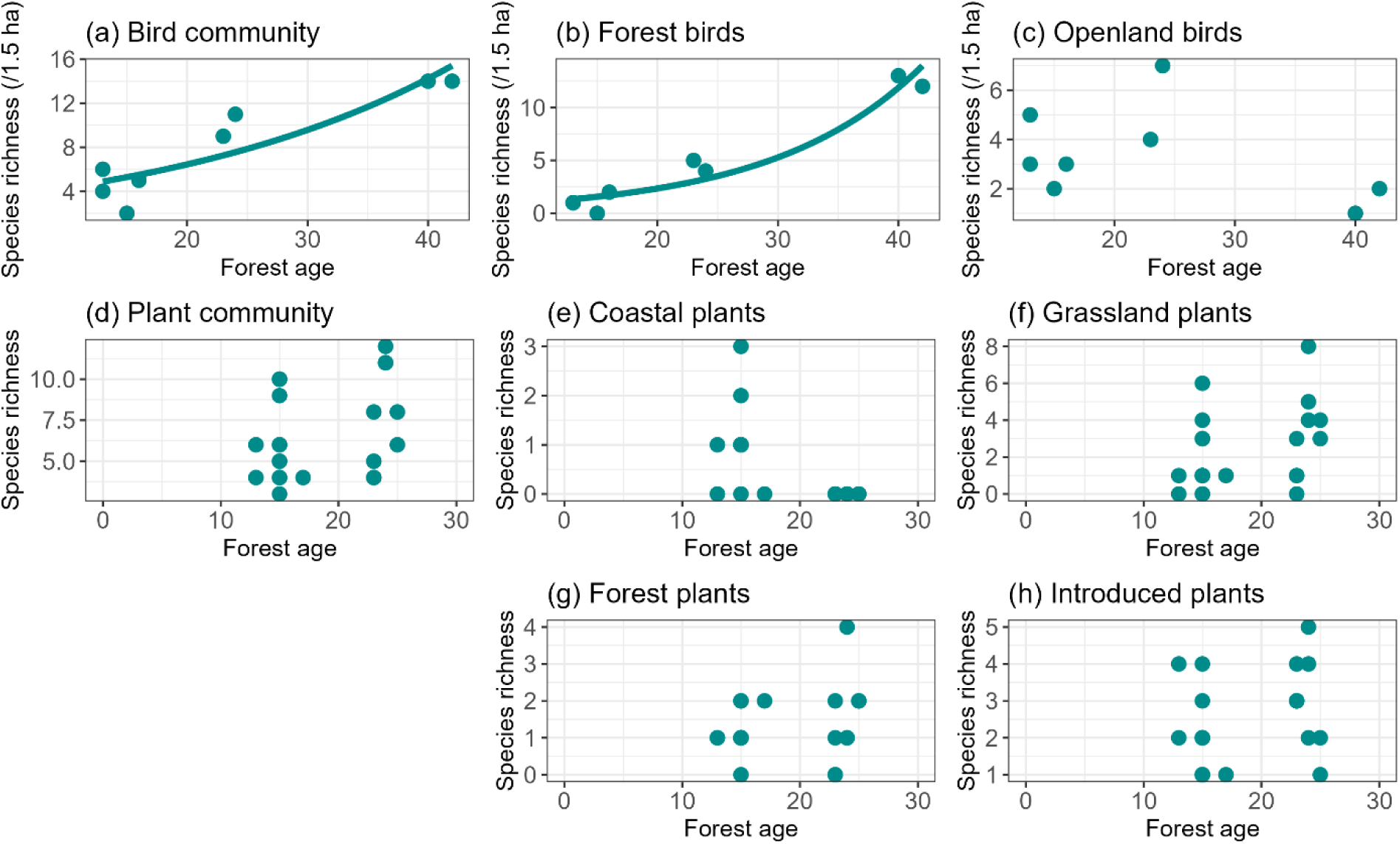
Relationships between bird (a-c) and plant (d-h) species richness and stand age. We excluded survey plots with stand age exceeding 60 years from this analysis. Predicted curves are shown only for statistically significant relationships. Dots indicate observed values.

## 4. Discussion

Although plantations in coastal ecosystems are far too pervasive, the extent to which plantation forests harbor coastal biodiversity has rarely been assessed or compared with that in natural coastal forests (Bonari *et al*. 2017; Muñoz-Reinoso 2021). We addressed this knowledge gap and showed that plantation coastal forests can still harbor community-level bird species richness comparable to that of natural forests. Plantations still retain coastal zonation patterns in bird species richness: in both forest types, forest bird species richness and abundance increased with distance from the shoreline. These results contrast with previous meta-analysis showing biodiversity in plantation forests is generally lower than in primary forests (Bremer & Farley 2010; Wang *et al*. 2022). This pattern may reflect the intrinsically low tree diversity of natural coastal forests. Intense environmental stress such as salt spray restricts persistence to a small number of stress-tolerant species (Du & Hesp 2020), often forming nearly monospecific stands of natural forests with lower species richness (Hasegawa 1984; Muñoz-Reinoso 2021). In these generally low-diversity systems, the negative impacts of plantation on avian communities may be weaker than those in other forest types.

However, plantations were unlikely to serve as complete substitutes for natural coastal forests for plant species. Community-level plant species richness was lower in plantations, and zonation patterns of plant species richness at the functional group level differed significantly between natural and plantation forests. Plantations, or wind fences, can buffer coastal stresses such as salt spray, wind, or sandy sediments, thereby enabling forest plant species to extend seaward following plantation activities (Grafals-Soto 2012). Site preparation likely acted as disturbances when plantation establishment, possibly providing suitable regeneration sites for coastal and introduced species further landward (Šebesta *et al*. 2021). These contrasting functional group-level responses suggest that, although biodiversity in plantation forests has often been assessed at the community level (Bremer & Farley 2010; Paillet *et al*. 2010), the functional group approach can identify underlying processes governing biodiversity patterns within plantations and thus better guide conservation strategies, particularly in forest systems with large variation in disturbance regimes.

Protecting the remaining continuum of natural forests and grasslands would be a central conservation priority in coastal ecosystems. Forests have been extensively converted, especially in lowland regions (Curran *et al*. 2004; Kitazawa *et al*. 2022b), and grasslands are among the most transformed and least protected terrestrial systems (Hoekstra *et al*. 2005). Since salt spray, sand drift, and wind maintain grassland habitats with scattered trees (Martínez & Psuty 2007), natural coastal zonation allows coexistence of contrasting groups, including forest and grassland species. In our study area, natural coastal vegetation actually harbored bird species of conservation concern, which strongly associated with mixed forest–grassland environments. Coastal ecosystems may thus serve as refugia for declining species of conservation concern in regions where intensive inland land use has eliminated lowland forests and grasslands (Acosta *et al*. 2009; Martínez & Psuty 2007; Sperandii *et al*. 2021).

While protecting existing natural forests and grasslands remains the highest conservation priority, plantation forests along the Ishikari coast also provided important habitat for Brown Shrike, a severely endangered openland bird species. Approximately 24% of the global Brown Shrike population breeds in this area (Kitazawa *et al*. 2022a), and nearly 90% of the individuals we observed were recorded in plantation forests. Existing studies suggest that scattered shrubs or trees in open habitats provide critical nesting and foraging habitat for both common and threatened openland birds (Fartmann *et al*. 2022; McLachlan 1991), and our results indicate that young plantation stands with moderate tree cover may play a similar role along the Ishikari coast (Fig.S4-S8, Table S2). These findings suggest that appropriate management in coastal plantations, such as retaining young, structurally open stands, could help buffer ongoing declines of the endangered species.

To minimize the negative impacts of plantations on coastal ecosystems, using natural zonation patterns as a reference can provide important management baselines. For example, pronounced differences in functional group-level plant species richness between sand-dune grasslands and plantation forests suggest that converting grasslands into plantations can substantially alter coastal ecosystems. Therefore, plantation activities are best confined to areas landward of coastal grasslands to avoid degradation and tree encroachment in grassland.

Intensive site preparation can facilitate the establishment and subsequent spread of neophytes into adjacent natural vegetation, and thus planting with no or less intensive site preparation may be essential for safeguarding native communities. Maintaining older stands is likely to benefit forest bird species, whereas unique coastal grassland ecosystems could still be conserved within plantation forests by retaining younger stands with sparse planting and low canopy height (Kawamura *et al*. 2025).

In this study, we provide, to our knowledge, the first comprehensive comparisons of biodiversity patterns between coastal plantations and natural forests and show that coastal plantations can harbor bird species richness comparable to that of natural forests at the community level. Our findings are consistent with earlier studies assessing biodiversity in old coastal plantations (Bonari *et al*. 2018) and further suggest that coastal plantations may offer, at least in part, opportunities to restore degraded lowland forest ecosystems. Afforestation in coastal ecosystems can also enhance disaster risk reduction, providing a means to simultaneously achieve biodiversity conservation and disaster mitigation (Iwachido *et al*. 2024). Care is needed when managing plantations in coastal ecosystems because stand age, although often a dominant driver explaining biodiversity in plantation forests (Hua *et al*. 2022), and community-level responses alone may not always serve as reliable management indicators. Our results suggest that taking distance from the shoreline (i.e., zonation patterns) into account and assessing functional group-level responses are also important in coastal systems, where species composition changes markedly along coastal disturbance gradients.

Although our results highlight the importance of protecting natural forests and grasslands, we may still underestimate the unique ecological functions of natural coastal forests, which are facing increasingly urgent threats from climate and land-cover change. Their regeneration dynamics are also likely to unfold over decadal timescales and, in this system, may depend on infrequent large disturbances such as eruptions (Hasegawa 1984; Saito & Higashi 1971). Future work should therefore evaluate longer-term dynamics and the responses of invertebrates (Ueno *et al*. 2025), which are key prey resources for the threatened bird species considered in this study.

## Acknowledgments

We thank the Environmental Citizen Department of Ishikari City, the Ishikari Subprefectural Bureau of Hokkaido Prefecture and the Hokkaido Regional Forest Office for granting permission and providing support for our field surveys. We are also grateful to Emi Takahashi, Natsumi Imai, Shunsuke Hori and Prof. Masaoki Takagi for helpful discussions. This study was made possible by the generous support of 46 donors through the Bird Research Support Project, and we express our deepest gratitude to them.

## Author contributions

Conceptualization: M.K., D.A., M.S.; Methodology: M.K.; Investigation: M.K., S.I., D.A.; Funding acquisition: M.K., S.I., D.A., M.S.; Supervision: M.S.; Writing – original draft: M.K.; Writing – review & editing: All authors.

