## Appendix for "Can coastal plantation forests substitute for natural coastal forests as bird and plant habitat? A test across coastal zonation"

**Title**

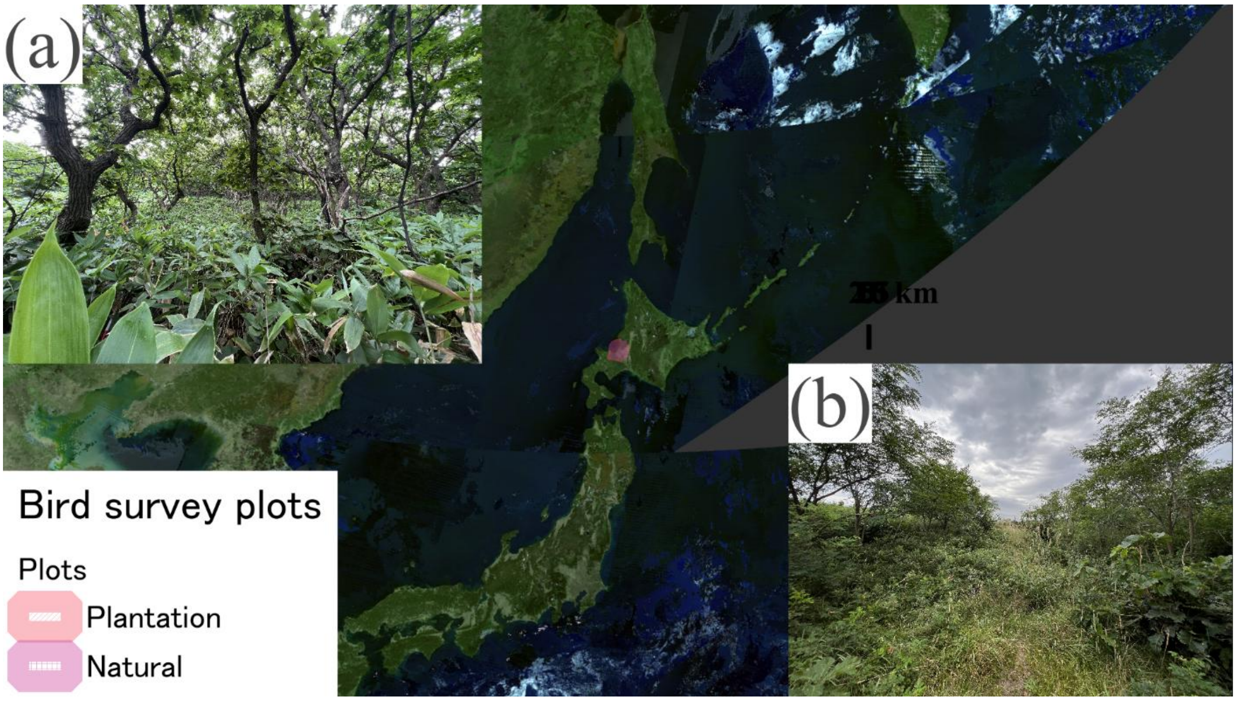


Figure S1. Distributions of bird survey plots. (a) Natural coastal forest composed of pure stands of *Quercus dentata*. (b) Young plantation coastal forest composed mainly of *Quercus dentata*.

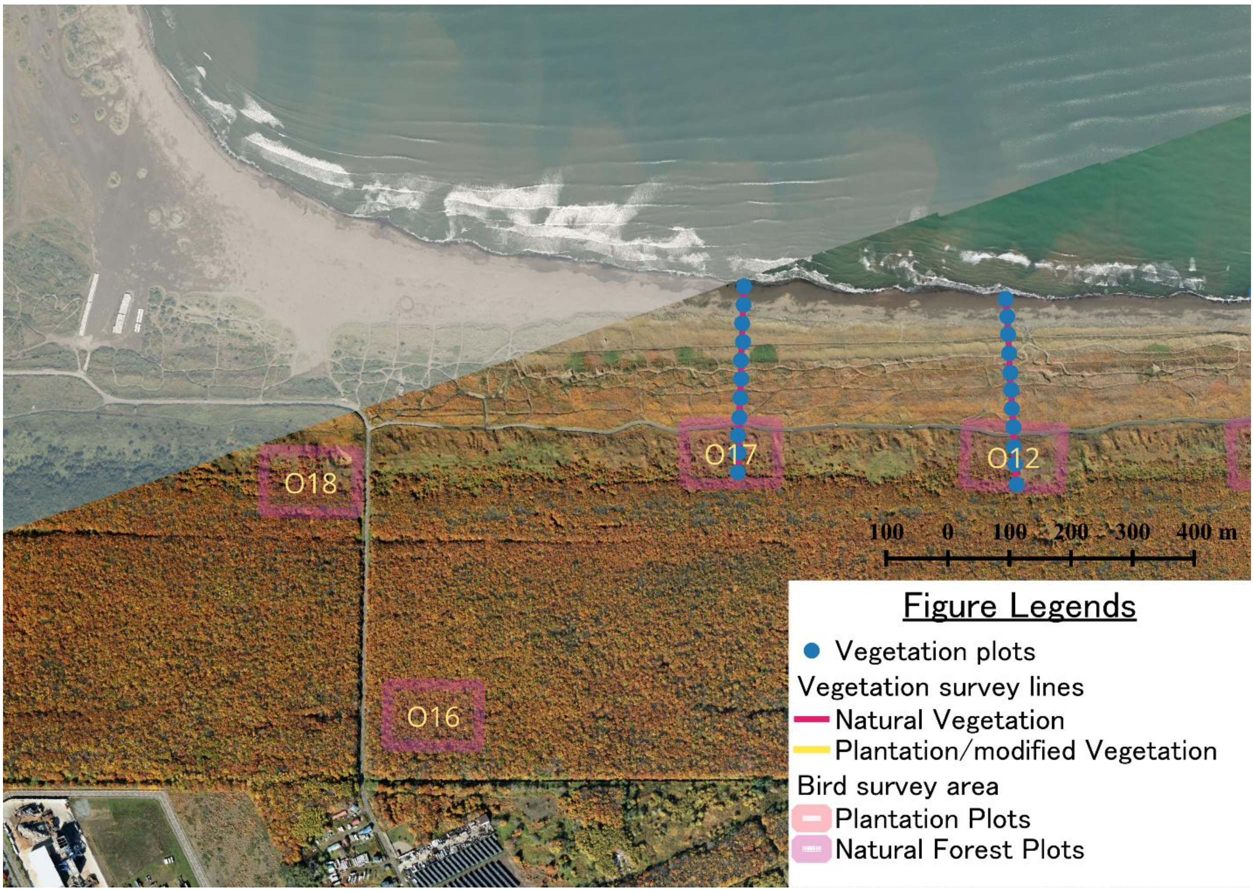


Figure S2. Distributions of survey plots in Ishikari coast. Blue plots indicate vegetation survey plots, and purple squares indicate bird survey plots. O12, O16, O17, O18 indicate bird survey plot IDs.

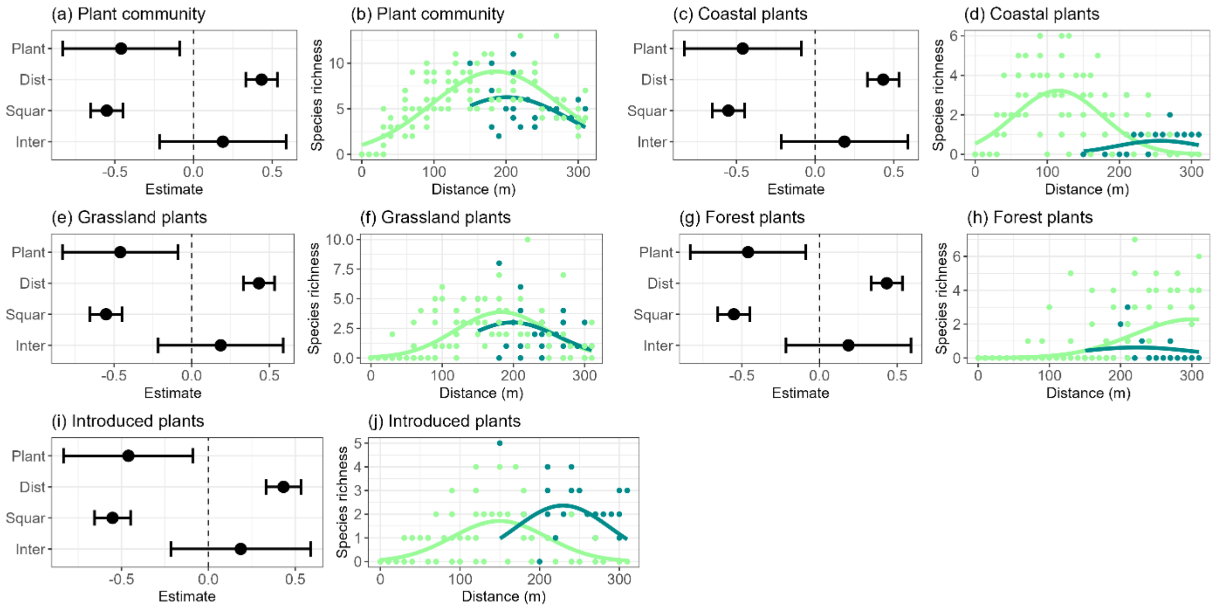


Figure S3. Relationships between plant community-level (a-b) and functional group-level (c-j) species richness and plantation category (Plant), distance from the shoreline (Dista), its squared term (Squar), and interaction between plantation category and distance (Inter) after excluding planted trees from dataset. Left-hand panel for each group shows estimated coefficients (dots) with 95% confidence intervals (error bars) for each predictor. Right-hand panels for each group show predicted relationships between distance from shoreline and species richness. Lines represent model predictions, and dots indicate observed values for natural forests (light green) and plantation forests (dark green).

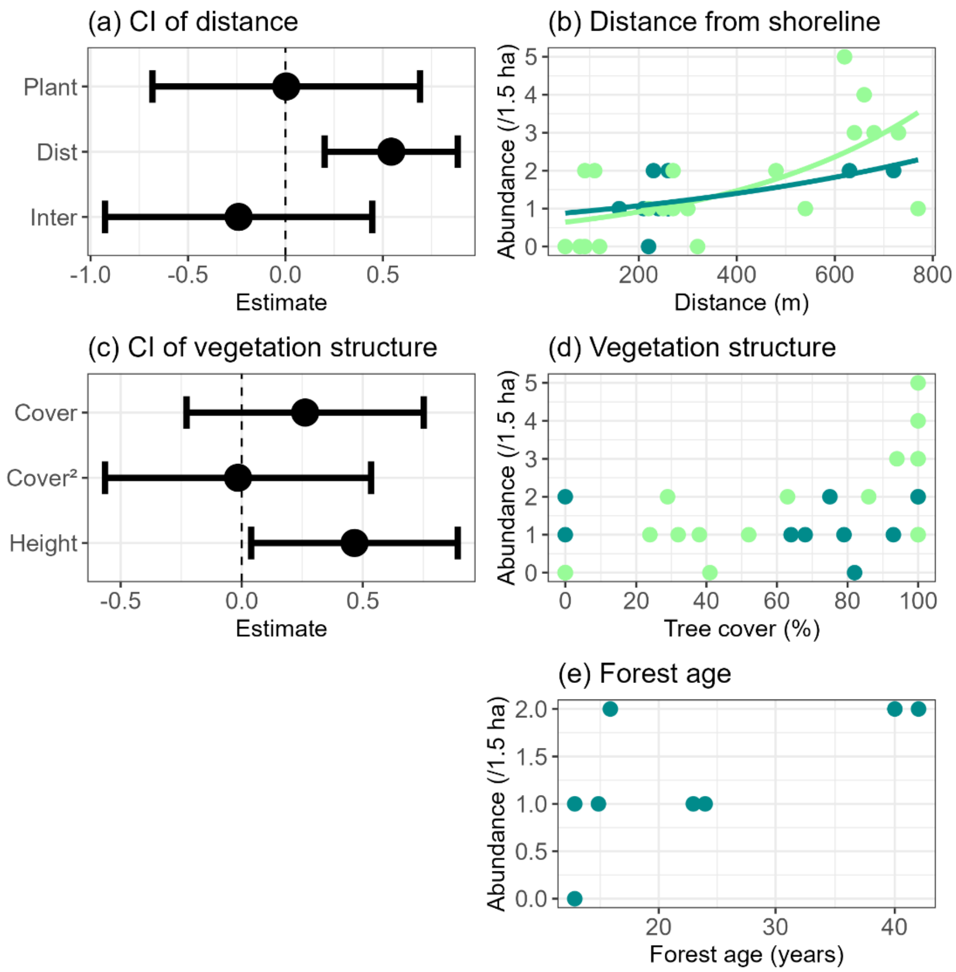


Figure S4. Relationships between Masked bunting abundance and environmental variables. (a)-(b) Estimated coefficients and confident intervals for plantation category (Plant), distance from the shoreline (Dist), and their interactions (Inter). (c)-(d) Estimated coefficients and confident intervals for wood cover (Cover) and its squared term ($Cover^2$), and vegetation height (Height). (e) Predicted relationships between Masked bunting abundance and stand age. Left-hand panels show estimated coefficients (dots) with 95% confidence intervals (error bars) for each predictor. Right-hand panels show predicted relationships between (b) distance from shoreline, (d) tree cover, and (e) stand age and Masked bunting abundance, respectively. Lines represent model predictions, and dots indicate observed values for natural forests (light green) and plantation forests (dark green).

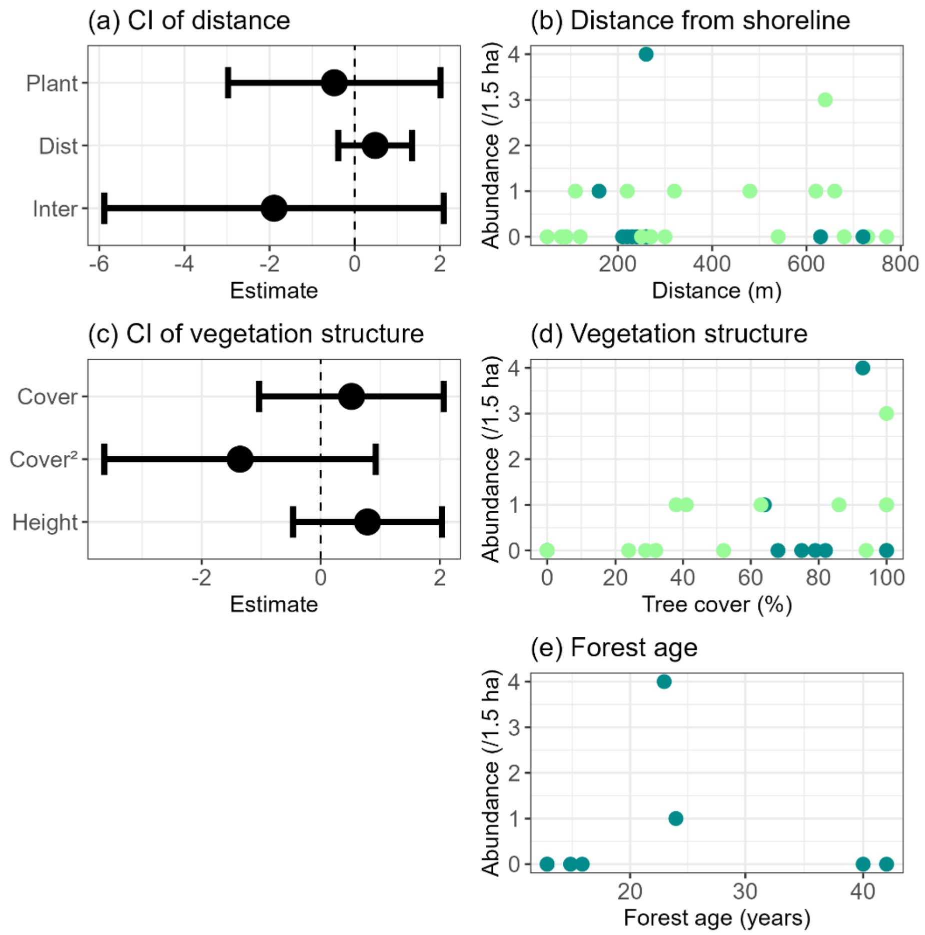


Figure S4. Relationships between Sakhalin grasshopper warbler abundance and environmental variables. (a)-(b) Estimated coefficients and confident intervals for plantation category (Plant), distance from the shoreline (Dist), and their interactions (Inter). (c)-(d) Estimated coefficients and confident intervals for wood cover (Cover) and its squared term ($Cover^2$), and vegetation height (Height). (e) Predicted relationships between Sakhalin grasshopper warbler abundance and stand age. Left-hand panels show estimated coefficients (dots) with 95% confidence intervals (error bars) for each predictor. Right-hand panels show predicted relationships between (b) distance from shoreline, (d) tree cover, and (e) stand age and Sakhalin grasshopper warbler abundance, respectively. Lines represent model predictions, and dots indicate observed values for natural forests (light green) and plantation forests (dark green).

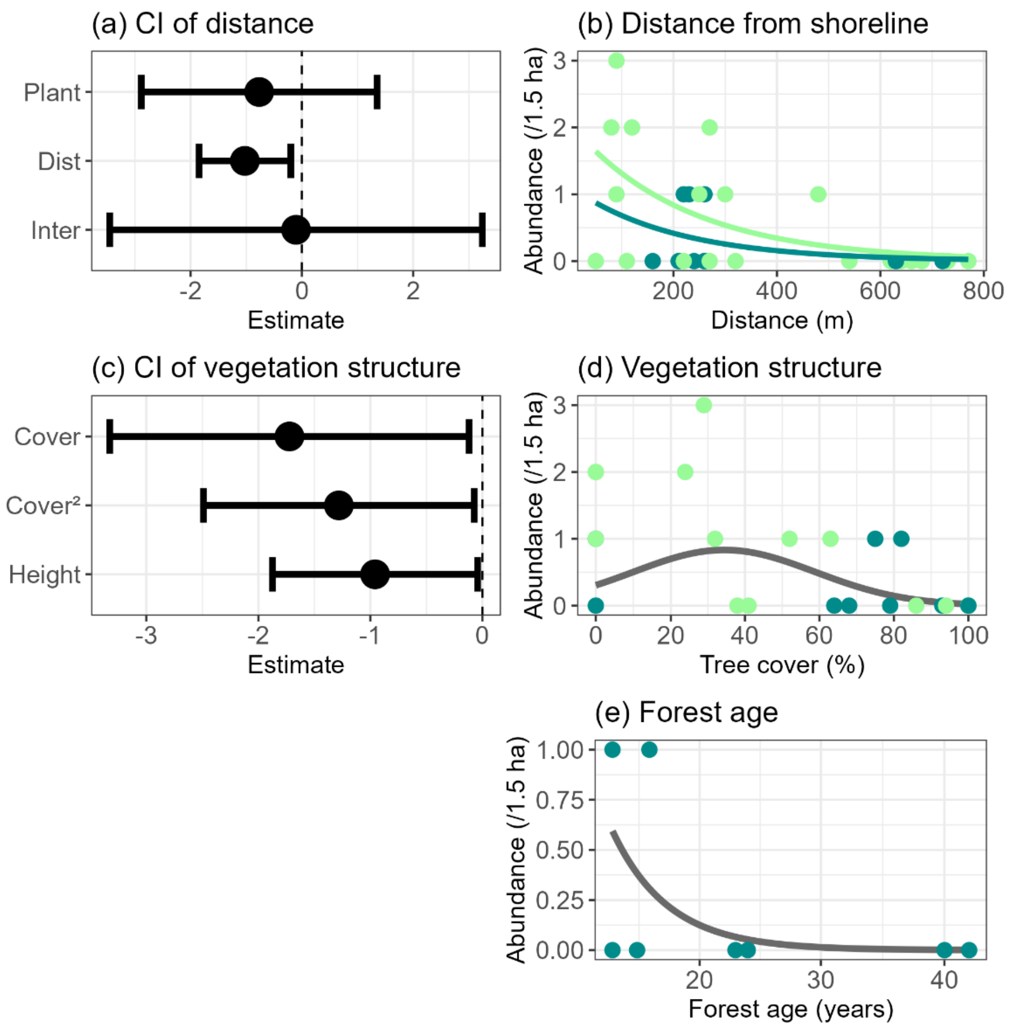


Figure S6. Relationships between Chestnut-eared bunting abundance and environmental variables. (a)-(b) Estimated coefficients and confident intervals for plantation category (Plant), distance from the shoreline (Dist), and their interactions (Inter). (c)-(d) Estimated coefficients and confident intervals for wood cover (Cover) and its squared term (Cover$^2$), and vegetation height (Height). (e) Predicted relationships between Chestnut-eared bunting abundance and stand age. Left-hand panels show estimated coefficients (dots) with 95% confidence intervals (error bars) for each predictor. Right-hand panels show predicted relationships between (b) distance from shoreline, (d) tree cover, and (e) stand age and Chestnut-eared bunting abundance, respectively. Lines represent model predictions, and dots indicate observed values for natural forests (light green) and plantation forests (dark green).

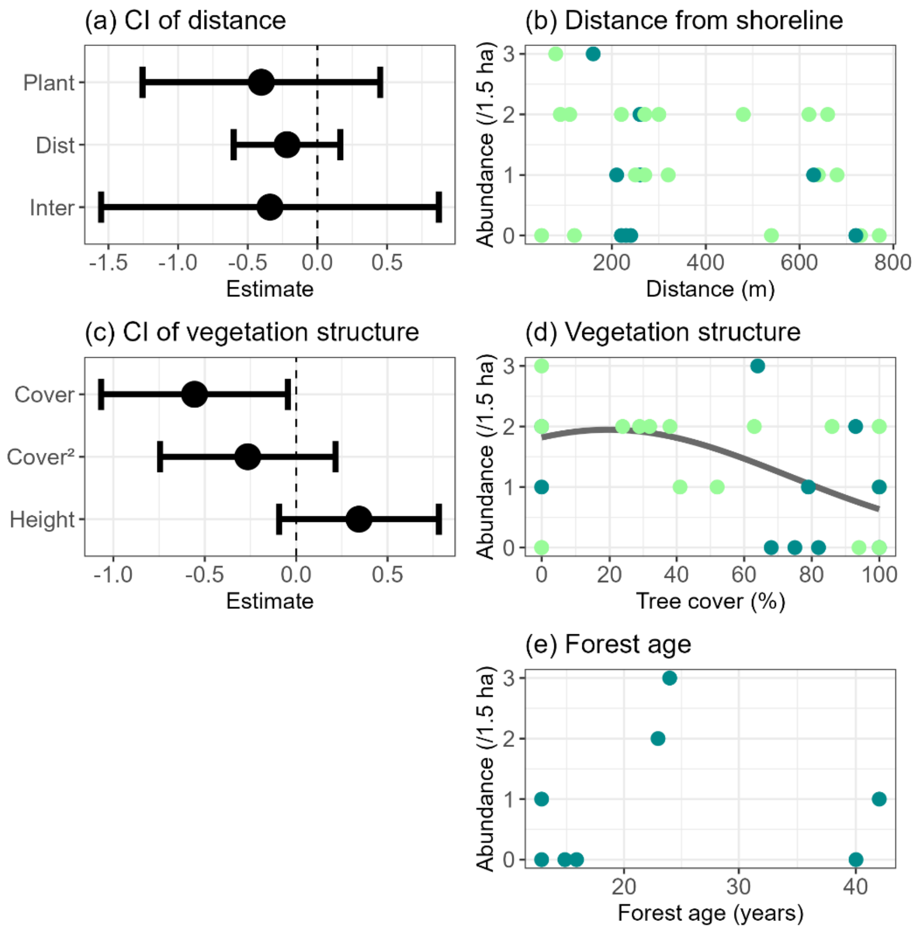


Figure S7. Relationships between Oriental greenfinch abundance and environmental variables. (a)-(b) Estimated coefficients and confident intervals for plantation category (Plant), distance from the shoreline (Dista), and their interactions (Inter). (c)-(d) Estimated coefficients and confident intervals for wood cover (Cover) and its squared term ($Cover^2$), and vegetation height (Height). (e) Predicted relationships between Oriental greenfinch abundance and stand age. Left-hand panels show estimated coefficients (dots) with 95% confidence intervals (error bars) for each predictor. Right-hand panels show predicted relationships between (b) distance from shoreline, (d) tree cover, and (e) stand age and Oriental greenfinch abundance, respectively. Lines represent model predictions, and dots indicate observed values for natural forests (light green) and plantation forests (dark green).

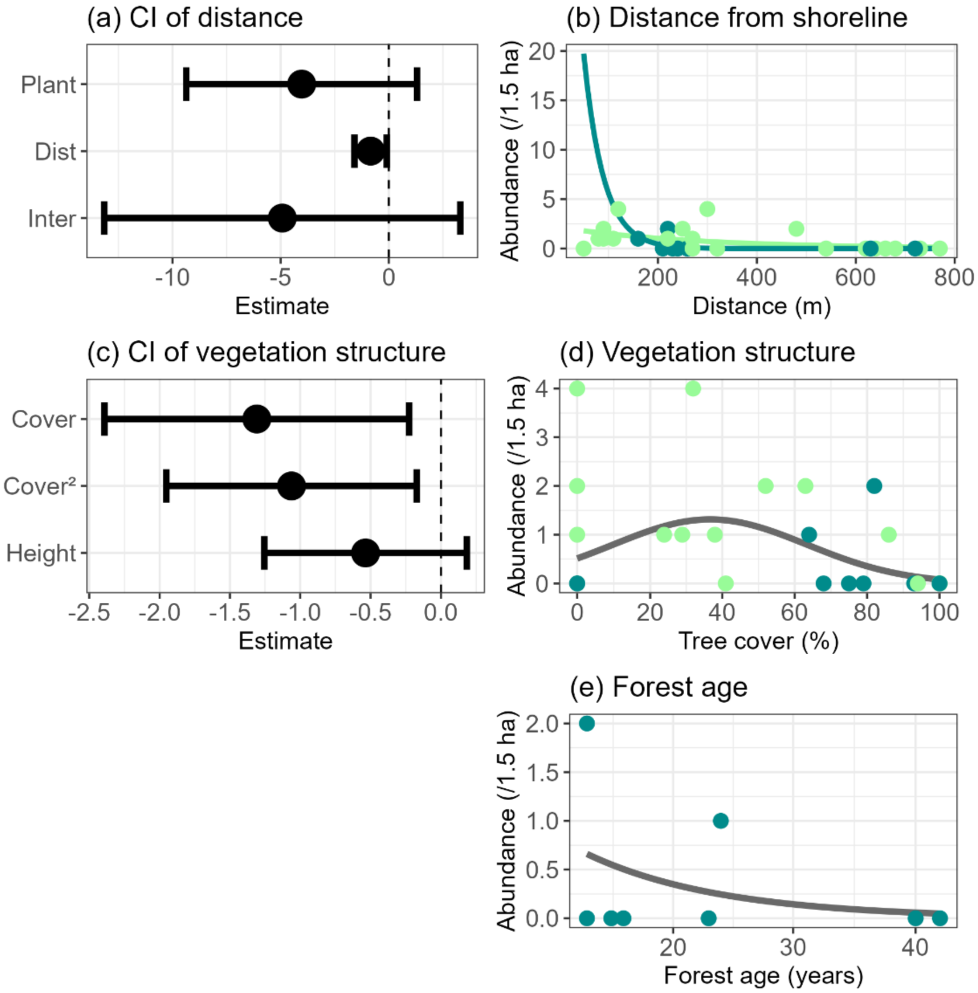


Figure S8. Relationships between Amur stonechat abundance and environmental variables. (a)-(b) Estimated coefficients and confident intervals for plantation category (Plant), distance from the shoreline (Dista), and their interactions (Inter). (c)-(d) Estimated coefficients and confident intervals for wood cover (Cover) and its squared term (Cover$^2$), and vegetation height (Height). (e) Predicted relationships between Amur stonechat abundance and stand age. Left-hand panels show estimated coefficients (dots) with 95% confidence intervals (error bars) for each predictor. Right-hand panels show predicted relationships between (b) distance from shoreline, (d) tree cover, and (e) stand age and Amur stonechat abundance, respectively. Lines represent model predictions, and dots indicate observed values for natural forests (light green) and plantation forests (dark green).

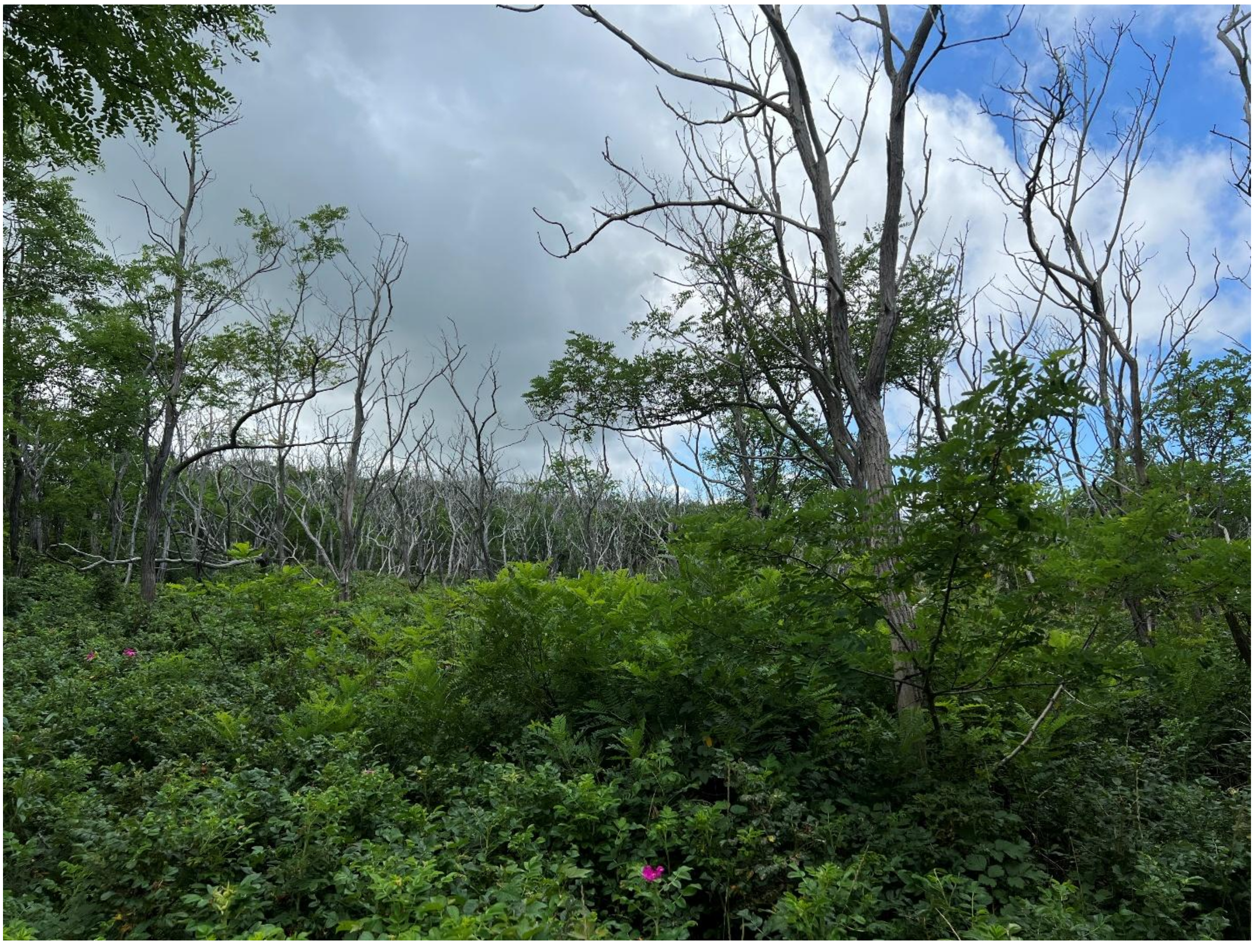

Figure S9. Photograph of a 61-year-old *Robinia pseudoacacia* plantation showing severe canopy dieback, with most canopy trees dead and the understorey dominated by *Rosa rugosa*.

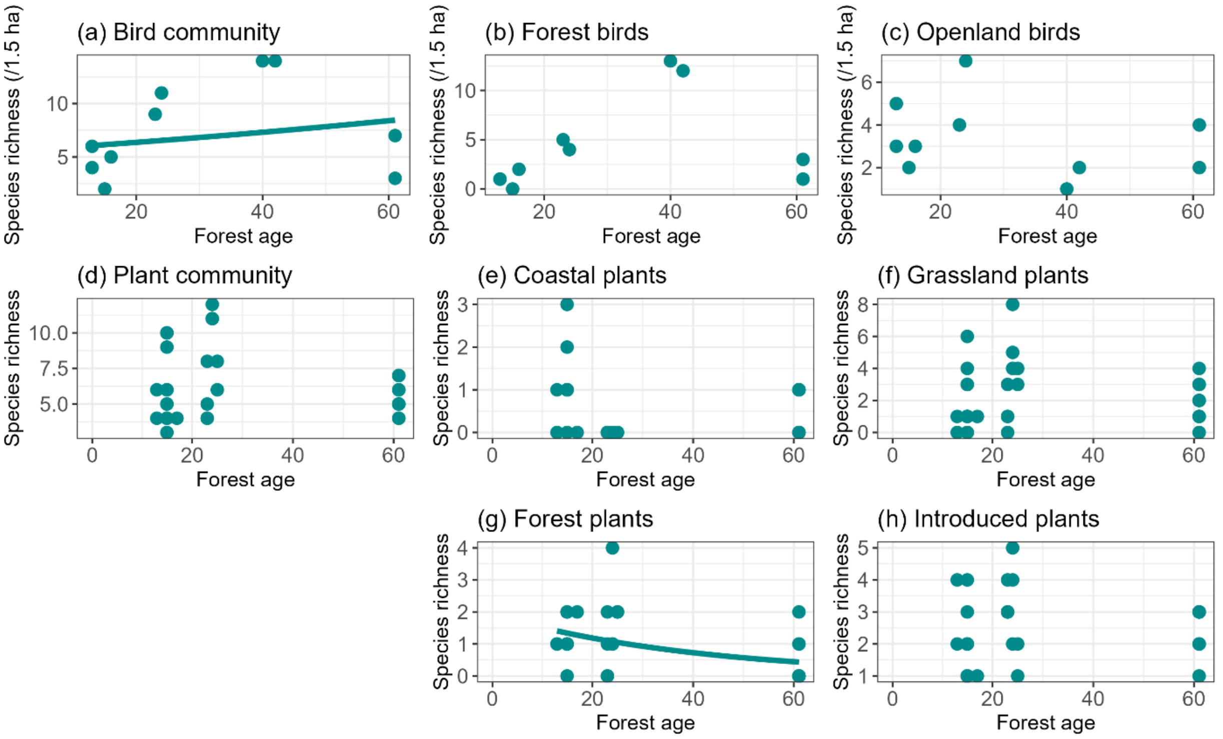


Figure. S10. Relationships between bird (a-c) and plant (d-h) species richness and stand age. We included survey plots with stand age exceeding 60 years in this analysis. Predicted curves are shown only for statistically significant relationships. Dots indicate observed values.

Table S1. Observed avian and vascular plant species. Detected individuals indicate the summed values of recorded bird abundance across all plots. Observed plots indicate the number of avian survey plots/vegetation quadrats that the species occurred.

| ID | Scientific name | Taxa | Functional group | Detected individuals | Observed plots |
|---|---|---|---|---|---|
| A01 | *Alauda arvensis* | Aves | Openland Species | 9 | 4 |
| A02 | *Motacilla alba* | Aves | Openland Species | 1 | 1 |
| A03 | *Acrocephalus orientalis* | Aves | Openland Species | 1 | 1 |
| A04 | *Emberiza fucata* | Aves | Openland Species | 16 | 11 |
| A05 | *Emberiza personata* | Aves | Openland Species | 45 | 24 |
| A06 | *Chloris sinica* | Aves | Openland Species | 36 | 21 |
| A07 | *Lanius cristatus* | Aves | Openland Species | 12 | 7 |
| A08 | *Lanius bucephalus* | Aves | Openland Species | 4 | 4 |
| A09 | *Saxicola stejnegeri* | Aves | Openland Species | 22 | 12 |
| A10 | *Emberiza schoeniclus* | Aves | Openland Species | 2 | 2 |
| A11 | *Locustella amnicola* | Aves | Openland Species | 14 | 9 |
| A12 | *Corvus corone* | Aves | Openland Species | 1 | 1 |
| A13 | *Calliope calliope* | Aves | Openland Species | 9 | 7 |
| A14 | *Acrocephalus bistrigiceps* | Aves | Openland Species | 1 | 1 |
| A15 | *Horornis diphone* | Aves | Forest Species | 20 | 16 |
| A16 | *Agropsar philippensis* | Aves | Forest Species | 8 | 6 |
| A17 | *Passer cinnamomeus* | Aves | Forest Species | 14 | 11 |
| A18 | *Streptopelia orientalis* | Aves | Forest Species | 16 | 15 |
| A19 | *Muscicapa dauurica* | Aves | Forest Species | 10 | 9 |
| A20 | *Ficedula narcissina* | Aves | Forest Species | 9 | 6 |
| A21 | *Poecile palustris* | Aves | Forest Species | 11 | 7 |
| A22 | *Phylloscopus borealoides* | Aves | Forest Species | 1 | 1 |
| A23 | *Phylloscopus coronatus* | Aves | Forest Species | 12 | 8 |
| A24 | *Zosterops japonicus* | Aves | Forest Species | 10 | 8 |
| A25 | *Turdus chrysolaus* | Aves | Forest Species | 11 | 10 |
| A26 | *Parus cinereus* | Aves | Forest Species | 11 | 9 |
| A27 | *Hypsipetes amaurotis* | Aves | Forest Species | 10 | 6 |
| A28 | *Dendrocopos major* | Aves | Forest Species | 7 | 7 |
| A29 | *Turdus cardis* | Aves | Forest Species | 5 | 5 |
| A30 | *Corvus macrorhynchos* | Aves | Forest Species | 2 | 1 |
| A31 | *Cuculus optatus* | Aves | Forest Species | 1 | 1 |
| A32 | *Urosphena squameiceps* | Aves | Forest Species | 4 | 3 |
| A33 | *Eophona personata* | Aves | Forest Species | 2 | 1 |
| A34 | *Sittiparus varius* | Aves | Forest Species | 1 | 1 |
| T01 | *Rosa rugosa* | Tracheophyta | Coastal Species | NA | 47 |
| T02 | *Leymus mollis* | Tracheophyta | Coastal Species | NA | 54 |
| T03 | *Linaria japonica* | Tracheophyta | Coastal Species | NA | 8 |
| T04 | *Carex kobomugi* | Tracheophyta | Coastal Species | NA | 30 |
| T05 | *Carex pumila* | Tracheophyta | Coastal Species | NA | 10 |
| T06 | *Artemisia stelleriana* | Tracheophyta | Coastal Species | NA | 2 |
| T07 | *Ixeris repens* | Tracheophyta | Coastal Species | NA | 23 |
| T08 | *Achillea alpina* | Tracheophyta | Coastal Species | NA | 7 |

| | | | | | |
|---|---|---|---|---|---|
| T09 | *Scutellaria strigillosa* | Tracheophyta | Coastal Species | NA | 4 |
| T10 | *Glehnia littoralis* | Tracheophyta | Coastal Species | NA | 11 |
| T11 | *Salsola komarovii* | Tracheophyta | Coastal Species | NA | 3 |
| T12 | *Calystegia soldanella* | Tracheophyta | Coastal Species | NA | 40 |
| T13 | *Lathyrus japonicus* | Tracheophyta | Coastal Species | NA | 7 |
| T14 | *Lonicera morrowii* | Tracheophyta | Grassland Species | NA | 6 |
| T15 | *Aralia elata* | Tracheophyta | Grassland Species | NA | 1 |
| T16 | *Rubus parvifolius* | Tracheophyta | Grassland Species | NA | 15 |
| T17 | *Elaeagnus umbellata* | Tracheophyta | Grassland Species | NA | 15 |
| T18 | *Ampelopsis glandulosa* | Tracheophyta | Grassland Species | NA | 4 |
| T19 | *Galium verum* | Tracheophyta | Grassland Species | NA | 3 |
| T20 | *Paederia foetida* | Tracheophyta | Grassland Species | NA | 2 |
| T21 | *Digitaria violascens* | Tracheophyta | Grassland Species | NA | 2 |
| T22 | *Miscanthus sinensis* | Tracheophyta | Grassland Species | NA | 57 |
| T23 | *Phalaris arundinacea* | Tracheophyta | Grassland Species | NA | 6 |
| T24 | *Luzula capitata* | Tracheophyta | Grassland Species | NA | 2 |
| T25 | *Eragrostis multicaulis* | Tracheophyta | Grassland Species | NA | 23 |
| T26 | *Deyeuxia epigeios* | Tracheophyta | Grassland Species | NA | 3 |
| T27 | *Carex caryophyllea* | Tracheophyta | Grassland Species | NA | 35 |
| T28 | *Carex miyabei* | Tracheophyta | Grassland Species | NA | 1 |
| T29 | *Artemisia montana* | Tracheophyta | Grassland Species | NA | 21 |
| T30 | *Artemisia japonica* | Tracheophyta | Grassland Species | NA | 4 |
| T31 | *Solidago virgaurea* | Tracheophyta | Grassland Species | NA | 4 |
| T32 | *Picris hieracioides* | Tracheophyta | Grassland Species | NA | 2 |
| T33 | *Aster ovatus* | Tracheophyta | Grassland Species | NA | 3 |
| T34 | *Hieracium umbellatum* | Tracheophyta | Grassland Species | NA | 4 |
| T35 | *Maianthemum dilatatum* | Tracheophyta | Grassland Species | NA | 2 |
| T36 | *Polygonatum humile* | Tracheophyta | Grassland Species | NA | 2 |
| T37 | *Thalictrum aquilegiifolium* | Tracheophyta | Grassland Species | NA | 1 |
| T38 | *Lysimachia vulgaris* | Tracheophyta | Grassland Species | NA | 1 |
| T39 | *Equisetum arvense* | Tracheophyta | Grassland Species | NA | 5 |
| T40 | *Pteridium aquilinum* | Tracheophyta | Grassland Species | NA | 4 |
| T41 | *Fallopia sachalinensis* | Tracheophyta | Grassland Species | NA | 1 |
| T42 | *Commelina communis* | Tracheophyta | Grassland Species | NA | 2 |
| T43 | *Dianthus superbus* | Tracheophyta | Grassland Species | NA | 7 |
| T44 | *Vicia japonica* | Tracheophyta | Grassland Species | NA | 23 |
| T45 | *Vicia cracca* | Tracheophyta | Grassland Species | NA | 1 |
| T46 | *Lathyrus palustris* | Tracheophyta | Grassland Species | NA | 1 |
| T47 | *Phragmites australis* | Tracheophyta | Grassland Species | NA | 3 |
| T48 | *Phragmites japonicus* | Tracheophyta | Grassland Species | NA | 1 |
| T49 | *Carex leucochlora* | Tracheophyta | Grassland Species | NA | 2 |
| T50 | *Equisetum hyemale* | Tracheophyta | Grassland Species | NA | 21 |
| T51 | *Solanum megacarpum* | Tracheophyta | Grassland Species | NA | 1 |
| T52 | *Quercus dentata* | Tracheophyta | Forest Species | NA | 45 |
| T53 | *Acer pictum* | Tracheophyta | Forest Species | NA | 2 |
| T54 | *Euonymus alatus* | Tracheophyta | Forest Species | NA | 15 |
| T55 | *Morus australis* | Tracheophyta | Forest Species | NA | 3 |
| T56 | *Betula platyphylla* | Tracheophyta | Forest Species | NA | 1 |
| T57 | *Vitis coignetiae* | Tracheophyta | Forest Species | NA | 12 |

| | | | | | |
|---|---|---|---|---|---|
| T58 | *Celastrus orbiculatus* | Tracheophyta | Forest Species | NA | 20 |
| T59 | *Euonymus sieboldianus* | Tracheophyta | Forest Species | NA | 1 |
| T60 | *Ligustrum obtusifolium* | Tracheophyta | Forest Species | NA | 3 |
| T61 | *Toxicodendron orientale* | Tracheophyta | Forest Species | NA | 9 |
| T62 | *Symplocos sawafutagi* | Tracheophyta | Forest Species | NA | 1 |
| T63 | *Maackia amurensis* | Tracheophyta | Forest Species | NA | 1 |
| T64 | *Galium odoratum* | Tracheophyta | Forest Species | NA | 5 |
| T65 | *Sasa palmata* | Tracheophyta | Forest Species | NA | 33 |
| T66 | *Pinus thunbergii* | Tracheophyta | Introduced Species | NA | 1 |
| T67 | *Populus alba* | Tracheophyta | Introduced Species | NA | 4 |
| T68 | *Robinia pseudoacacia* | Tracheophyta | Introduced Species | NA | 18 |
| T69 | *Amorpha fruticosa* | Tracheophyta | Introduced Species | NA | 16 |
| T70 | *Oenothera biennis* | Tracheophyta | Introduced Species | NA | 4 |
| T71 | *Cakile edentula* | Tracheophyta | Introduced Species | NA | 10 |
| T72 | *Dactylis glomerata* | Tracheophyta | Introduced Species | NA | 41 |
| T73 | *Festuca rubra* | Tracheophyta | Introduced Species | NA | 26 |
| T74 | *Phleum pratense* | Tracheophyta | Introduced Species | NA | 1 |
| T75 | *Anthoxanthum odoratum* | Tracheophyta | Introduced Species | NA | 8 |
| T76 | *Agrostis gigantea* | Tracheophyta | Introduced Species | NA | 3 |
| T77 | *Elymus repens* | Tracheophyta | Introduced Species | NA | 8 |
| T78 | *Lolium multiflorum* | Tracheophyta | Introduced Species | NA | 1 |
| T79 | *Plantago lanceolata* | Tracheophyta | Introduced Species | NA | 7 |
| T80 | *Hypochaeris radicata* | Tracheophyta | Introduced Species | NA | 18 |
| T81 | *Solidago gigantea* | Tracheophyta | Introduced Species | NA | 7 |
| T82 | *Erigeron canadensis* | Tracheophyta | Introduced Species | NA | 2 |
| T83 | *Cirsium arvense* | Tracheophyta | Introduced Species | NA | 1 |
| T84 | *Cota tinctoria* | Tracheophyta | Introduced Species | NA | 1 |
| T85 | *Rumex acetosella* | Tracheophyta | Introduced Species | NA | 4 |
| T86 | *Rumex obtusifolius* | Tracheophyta | Introduced Species | NA | 2 |

Table S2. Estimated effects of distance from shoreline, max tree height, and tree cover within bird survey plots on bird species richness. Separate GLMMs were fitted for each response and explanatory variable, with species richness at each level as the response variable.

| **Response variables** | **Explanatory variables** | **Coefficient** | **AIC** |
|---|---|---|---|
| Community-level species richness | Distance | 0.001 | 159.8 |
| Community-level species richness | Max tree height | 0.01 | 163.1 |
| Community-level species richness | Tree cover | 0.002 | 159.1 |
| Openland species richness | Distance | -0.001 | 114.8 |
| Openland species richness | Max tree height | -0.1 | 119.4 |
| Openland species richness | Tree cover | -0.005 | 118.8 |
| Forest species richness | Distance | 0.003 | 141.4 |
| Forest species richness | Max tree height | 0.3 | 147.4 |
| Forest species richness | Tree cover | 0.02 | 139.2 |
